# Co-recognition of histone acetylation and H3K4 trimethylation by GTE4-EML complex in Arabidopsis

**DOI:** 10.1101/2024.07.25.605061

**Authors:** Feng Qian, Qiang-Qiang Zhao, Jin-Xing Zhou, Dan-Yang Yuan, Yin-Na Su, Lin Li, She Chen, Xin-Jian He

**Affiliations:** National Institute of Biological Sciences, Beijing 102206, China; Tsinghua Institute of Multidisciplinary Biomedical Research, Tsinghua University, 100084, Beijing, China

**Keywords:** histone acetylation, H3K4me3, Bromodomain, GTE4, EML1, transcription, development

## Abstract

Although histone acetylation and H3K4 trimethylation (H3K4me3) are well-known histone marks associated with active transcription, how they cooperate to regulate transcription remains largely unclear in plants. Our study revealed that the Bromodomain and Extra-terminal (BET) protein GTE4 binds to acetylated histone and forms a complex with the redundant H3K4me3-binding EMSY-Likeproteins EML1 or EML2 (EML1/2) in *Arabidopsis thaliana*. The *eml1 eml2* (*eml1/2*) double mutant exhibited a morphological phenotype similar to the *gte4* mutant, and most of the *gte4*-mediated differentially expressed genes were co-regulated in the *eml1/2* mutant. Through chromatin immunoprecipitation followed by deep sequencing (ChIP-seq), we found that GTE4 and EML2 co-occupy protein-coding genes enriched with both histone acetylation and H3K4me3, exhibiting a synergistic effect on the association of the GTE4-EML complex with chromatin. The association of GTE4 with chromatin requires both the Bromodomain and the EML-interacting domain. This study identified a previously uncharacterized complex and uncovered how it cooperatively recognizes histone acetylation and H3K4me3 to facilitate gene transcription at the whole-genome level in Arabidopsis.

## Introduction

Histone modifications play a crucial role in regulating the action of chromatin, including gene transcription, DNA replication, and DNA damage repair (Karlic et al., 2010; Bar-Ziv et al., 2016; Clouaire et al., 2018). One well-studied modification is histone acetylation at lysine sites, which is associated with increased chromatin accessibility and transcriptional activation (Struhl, 1998; Bannister and Kouzarides, 2011; Berr et al., 2011). Histone acetylation can reduce the interaction between histones and DNA by neutralizing the positive charge of lysine sites, thereby mediating transcriptional activation (Andrews et al., 2016). Moreover, the conserved bromodomain was identified as a “reader” of histone acetylation to activate transcription (Marmorstein and Zhou, 2014). Histone methylation at lysine sites is another histone modification type that is associated with transcription. The locations of methylated lysine sites are crucial for determining the effects of histone methylation on transcription. While the H3 lysine 4 and 36 methylation is associated with actively transcribed genes, the H3 lysine 9 and 27 methylation is associated with transcriptional repression (Suganuma and Workman, 2011; Liu et al., 2018). Especially, histone H3K4 trimethylation (H3K4me3) is a histone mark that is associated with transcriptional activation (Zhang et al., 2009; Fromm and Avramova, 2014; Jambhekar et al., 2019). A subset of PHD fingers were identified as H3K4me3 “readers” to mediate the connection between H3K4me3 and transcription (Berr et al., 2011; Sanchez and Zhou, 2011). Although both histone acetylation and H3K4me3 are known to associate with transcriptional activation, it is largely unclear how they cooperate to regulate transcription at the whole-genome level in plants.

The bromodomain is present in various chromatin-related proteins, including histone acetyltransferases, chromatin remodeling complex components, and transcriptional regulators in eukaryotes (Pandey et al., 2002; Zeng and Zhou, 2002; Filippakopoulos et al., 2012). In *Arabidopsis thaliana*, twenty-nine bromodomain proteins have been identified (Pandey et al., 2002). However, only a small part of them has been characterized as histone acetylation “readers” (Zhang et al., 2016; Nie et al., 2019; Potok et al., 2019; Sijacic et al., 2019; Luo et al., 2020; Jaronczyk et al., 2021; Yu et al., 2021; Guo et al., 2022). Bromodomain and extra-terminal domain (BET) proteins constitute a significant subfamily of bromodomain proteins in eukaryotes (Pandey et al., 2002; Zeng and Zhou, 2002). In fungi and metazoans, BET proteins have been shown to bind to acetylated histones, thereby regulating transcription (Ladurner et al., 2003; Wang et al., 2013), DNA damage signaling (Floyd et al., 2013), and cell cycling (LeRoy et al., 2008). In Arabidopsis, there are 12 BET proteins collectively known as GTE1-GTE12 (GLOBAL TRANSCRIPTION FACTOR GROUP E 1-12).

GTE1/IMB1 is involved in seed germination by facilitating the activation of genes related to cell wall metabolism and plastid-encoded genes (Duque and Chua, 2003). GTE6 mediates the transcriptional activation of *AS1*, which encodes a MYB-domain protein involved in leaf development (Chua et al., 2005). GTE4 is essential for proper root and leaf development as well as plant immunity (Airoldi et al., 2010; Zhou et al., 2022). Additionally, GTE9 and GTE11, two closely related BET proteins, interact with BT2 (BTB AND TAZ DOMAIN PROTEIN 2) to mediate responses to ABA and sugar signaling (Misra et al., 2018). Despite the reported biological functions of these Arabidopsis BET proteins, the specific involvement of the bromodomain in their functions in plants has yet to be determined.

Yeast and metazoan BET proteins typically possess two to five tandem bromodomains that specifically bind to different combinations of acetylated lysine sites on histones, enabling the recognition of specific chromatin regions (Dyson et al., 2001; Pandey et al., 2002; Taniguchi, 2016). The tandem arrangement of bromodomains enhances their binding affinity to acetylated histones synergistically (Moriniere et al., 2009; Filippakopoulos et al., 2012). In contrast, all Arabidopsis BET proteins have a single bromodomain (Pandey et al., 2002). Previous studies identified GTE1 as an acetylated-histone-binding protein through *in vitro* assays, indicating that Arabidopsis BET family proteins likely possess a conserved ability with acetylated histones (Zhao et al., 2018). However, it remains unclear how these single-bromodomain BET proteins discriminate between chromatin regions with varying types or levels of histone acetylation. It is also unknown whether unique mechanisms for chromatin recognition have evolved in plants to compensate for the absence of tandem-bromodomain BET proteins. Furthermore, while several BET proteins in yeast and metazoans have been shown to interact with other chromatin regulators, such interactions have not yet been identified and characterized in plants (Floyd et al., 2013; Wang et al., 2013; Taniguchi, 2016). Therefore, it is necessary to identify and study potential BET-interacting proteins in plants.

In this study, we identified five closely related Arabidopsis BET proteins (GTE2, GTE3, GTE4, GTE5, and GTE7) as acetylated-histone binding proteins and demonstrated that the binding of GTE4 to acetylated histone facilitates the association of GTE4 with chromatin. Moreover, we found that GTE4 interacts with two redundant single-Tudor domain proteins: EML1 and EML2 (EML1/2). The single-Tudor domain of EML1 was previously identified as a plant-specific “reader” of histone H3K4 trimethylation (H3K4me3) (Zhao et al., 2018). We demonstrated that GTE4 and EML1/2 co-occupy chromatin regions that are enriched with both histone acetylation and H3K4me3, and the occupancy of GTE4 and EML1/2 on chromatin is mutually dependent. Moreover, we have identified a conserved α-helix domain in GTE4 that is responsible for interacting with EML1 and EML2. These results provide insights into the cooperative recognition mechanism employed by single-bromodomain BET proteins and single-Tudor domain proteins and thereby shed light on the plant-specific combinatorial recognition of histone acetylation and H3K4me3.

## Results

### Identification and characterization of the GTE4 bromodomain

To identify acetylated histone-binding proteins in Arabidopsis, we performed a pull-down assay by using the histone H4 peptide with acetylation at lysine 5, 8, 12, and 16 (H4K5/8/12/16ac) as a bait and then detected acetylated histone-interacting proteins by affinity purification followed by mass spectrometry (AP-MS). Through this analysis, we identified five bromodomain and extra terminal domain (BET) proteins (GTE2, GTE3, GTE4, GTE5, and GTE7) as acetylated histone-interacting proteins (Figure 1A; Dataset 1). While GTE1 was not identified as an acetylated-histone-binding BET protein in our current AP-MS analysis, a previous *in vitro* pull-down assay demonstrated its strong binding affinity with the histone H3K5ac peptide (Zhao et al., 2018). Therefore, the acetylated-histone-binding ability is likely to be conserved for all the Arabidopsis BET proteins.

**Figure 1.**
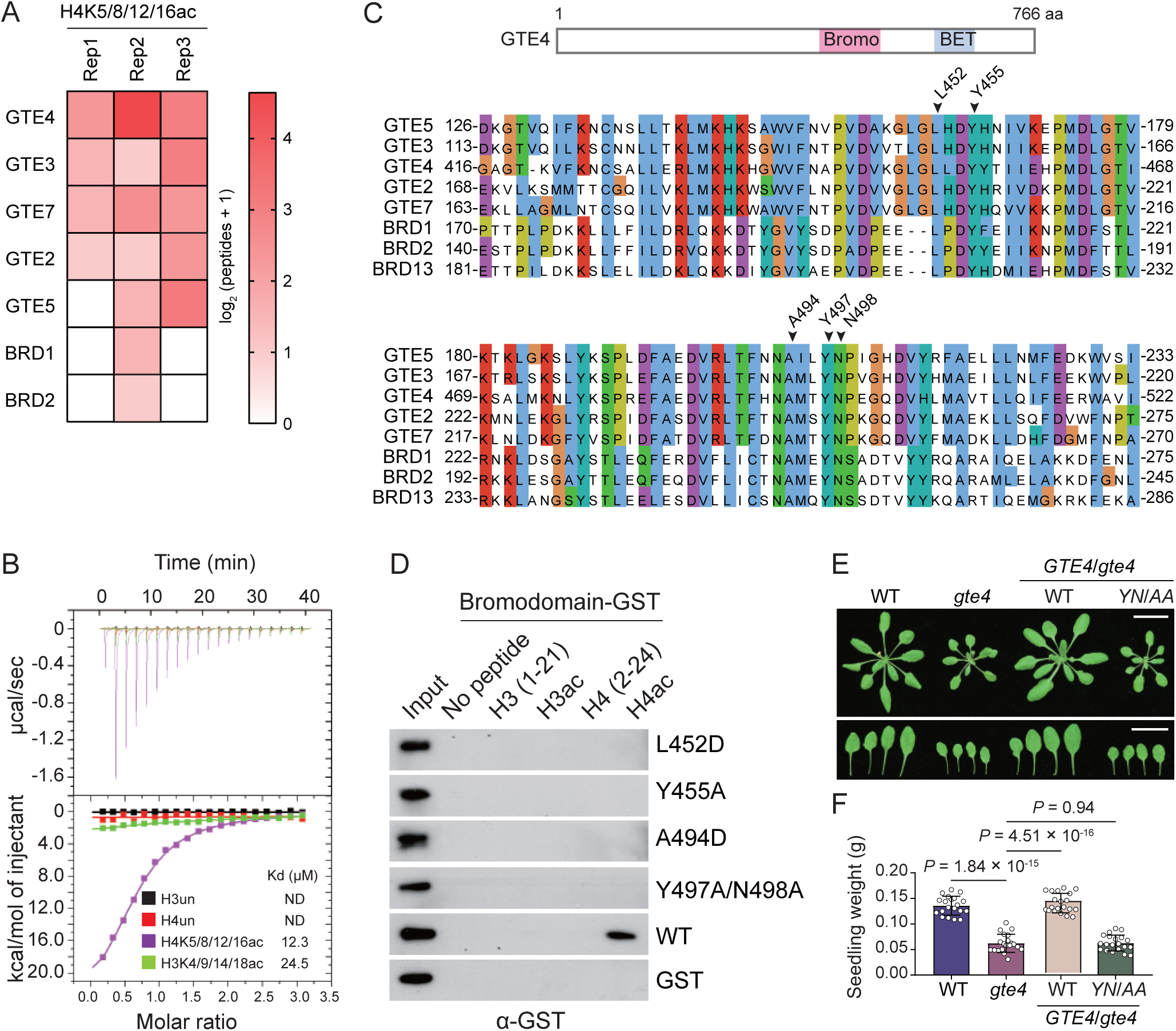
GTE4 and its homologs as major readers of histone acetylation in Arabidopsis plants. (A) Identification of GTEs as acetylated-histone-binding proteins by AP-MS. Acetylated-histone binding proteins were identified by AP-MS using the histone H4 peptide with acetylation of lysine 5, 8, 12, and 16 (H4K5/8/12/16ac) as a bait. (B) Determination of the binding affinity of the GTE4 bromodomain with histone peptides by ITC assays. The histone peptides used in ITC include H3 (1-19 aa), H3K4/9/14/18ac (1-26 aa), H4 (1-20 aa), and H4K5/8/12/16ac (1-20 aa). (C) Analyses of the bromodomains in GTE proteins. The schematic diagram of conserved domains of GTE4 is shown in the upper panel, and the alignment of the bromodomains in indicated GTE and BRD proteins is shown in the lower panel. The conserved residues mutated in this study are labeled on the top of aligned sequences. (D) Determination of the binding of wild-type and mutated bromodomains of GTE4 to histone peptides with and without lysine acetylation. Biotin-labeled histone peptides were incubated with wild-type and mutated bromodomains of GTE4 and subjected to pull-down assays. H3ac and H4ac represent H3K4/9/14/18ac (1-26 aa) and H4K5/8/12/16ac (1-18 aa), respectively. (E,F) Effect of the bromodomain mutations on the function of GTE4 in Arabidopsis plants. Wild-type *GTE4* and mutated *GTE4-Y497A/N498A* transgenes were transformed into the *gte4* mutant for complementation testing. The morphological phenotype of 21-day-old-plants, including the 3^rd^, 4^th^, 5^th^, and 6^th^ rosette leaves (E), and the statistical analysis of 21-day-old seedling weight (F) are shown. The values of plant weight of 21-day-old seedlings are shown as means ± SD (n = 20). P values were determined by Student t test with two tails. Bar, 2 cm.

Among the acetylated-histone-binding BET proteins identified by our AP-MS analysis, GTE4 has been extensively studied. A previous study has shown that GTE4 is involved in regulating root and leaf development (Airoldi et al., 2010). To investigate whether and how the binding of GTE4 to acetylated histone contributes to the function of GTE4, we focused our current study on GTE4. We expressed and purified the bromodomain of GTE4, which was tagged with GST, from *E. coli*. Subsequently, we conducted an ITC (isothermal titration calorimetry) assay to determine the binding affinity of the GTE4 bromodomain for acetylated histones. The ITC assay indicated that the bromodomain of GTE4 displayed a strong affinity for the histone H4 peptide with acetylation at lysine 5, 8, 12, and 16 (H4K5/8/12/16ac) and a low binding affinity for the H3 histone with acetylation at lysine 4, 9, 14, and 18 (H3K4/9/14/18ac), but did not exhibit a detectable binding affinity for unmodified H3 or H4 histone peptides (Figure 1B). A histone peptide pull-down assay further supported these findings, demonstrating that the bromodomain of GTE4 exhibits a high binding affinity for H4K5/8/12/16ac and to a lesser extent for H3K4/9/14/18ac (Supplemental Figure 1). These results establish that the bromodomain of GTE4 has a specific binding affinity for acetylated histone.

The bromodomain, found in BRD1, BRD2, and BRD13, which are subunits of BRM-associated SWI/SNF complexes, shares a conserved ability to bind acetylated histones (Zhao et al., 2018; Yu et al., 2021; Guo et al., 2022). Through sequence alignment, we observed that the bromodomain of GTE4 contains residues L452, Y455, A494, Y497, and N498 that are conserved not only in BET proteins but also in BRD1, BRD2, and BRD13 (Figure 1C). To investigate the impact of these conserved residues on the binding of the GTE4 bromodomain to acetylated histones, we introduced point mutations (L452D, Y455A, A494D, and Y497A/N498A) and conducted a histone peptide pull-down assay. The results of the pull-down assay revealed that the GTE4 bromodomain shows a high binding affinity for H4K5/8/12/16ac but not for unmodified H4 (Figure 1D; Supplemental Figure 1). Additionally, the pull-down assay demonstrated that all the introduced mutations completely disrupt the binding of the GTE4 bromodomain to acetylated H4 (Figure 1D). These findings strongly suggest that the conserved residues in the GTE4 bromodomain are required for binding to acetylated histone.

Furthermore, we introduced the *Y497A/N498A* mutation into the *GTE4* transgene and then determined whether the conserved bromodomain of GTE4 is required for the biological function of GTE4 in Arabidopsis plants. Consistent with the previous report on the *gte4* mutant (Airoldi et al., 2010), our study indicated that the *gte4* mutant (SALK_083697) exhibited smaller rosette size, reduced seedling weight, and jagged leaves compared to the wild type (Figure 1E, 1F). Complementation testing demonstrated that the developmental defects in the *gte4* mutant were restored by the wild-type *GTE4* transgene but not by the *GTE4* transgene carrying the *Y497A/N498A* mutation (Figure 1E, 1F; Supplemental Figure 2). These findings indicate that the conserved bromodomain is crucial for the biological function of GTE4 in Arabidopsis plants.

### GTE4 interacts with the H3K4me3-binding homologs EML1 and EML2

To investigate the mechanisms underlying the regulation of chromatin state by GTE4, we generated transgenic Arabidopsis plants expressing GTE4 tagged by a FLAG epitope (GTE4-FLAG) and then identified GTE4-interacting proteins by AP-MS using anti-FLAG antibody. Our AP-MS result revealed that two closely related EMSY-LIKE (EML) proteins, EML1 and EML2 (EML1/2), were co-purified with GTE4, indicating a potential interaction between GTE4 and EML1/2 (Figure 2A; Dataset 2). To further investigate this interaction, we generated transgenic Arabidopsis plants expressing FLAG-tagged EML1/2. Subsequently, we performed AP-MS to isolate proteins that interact with EML1/2. Interestingly, we found that GTE4 was among the proteins co-purified with EML1/2 (Figure 2A; Dataset 2). EML1/2 proteins possess an Agenet/Tudor domain and an EMSY N-terminal (ENT) domain (Tsuchiya and Eulgem, 2011). A previous study has demonstrated that the Agenet domain of EML1 can bind to trimethyl H3K4 (H3K4me3) peptide (Zhao et al., 2018), and the specificity of this binding was confirmed using histone peptide arrays and pull-down assays in our current study (Supplemental Figure 3A, 3B). Considering that both histone acetylation and H3K4me3 are associated with active transcription (Struhl, 1998; Bannister and Kouzarides, 2011), the co-recognition of GTE4-EML1 is most likely to regulate transcription at chromatin regions enriched with both histone acetylation and H3K4me3.

**Figure 2.**
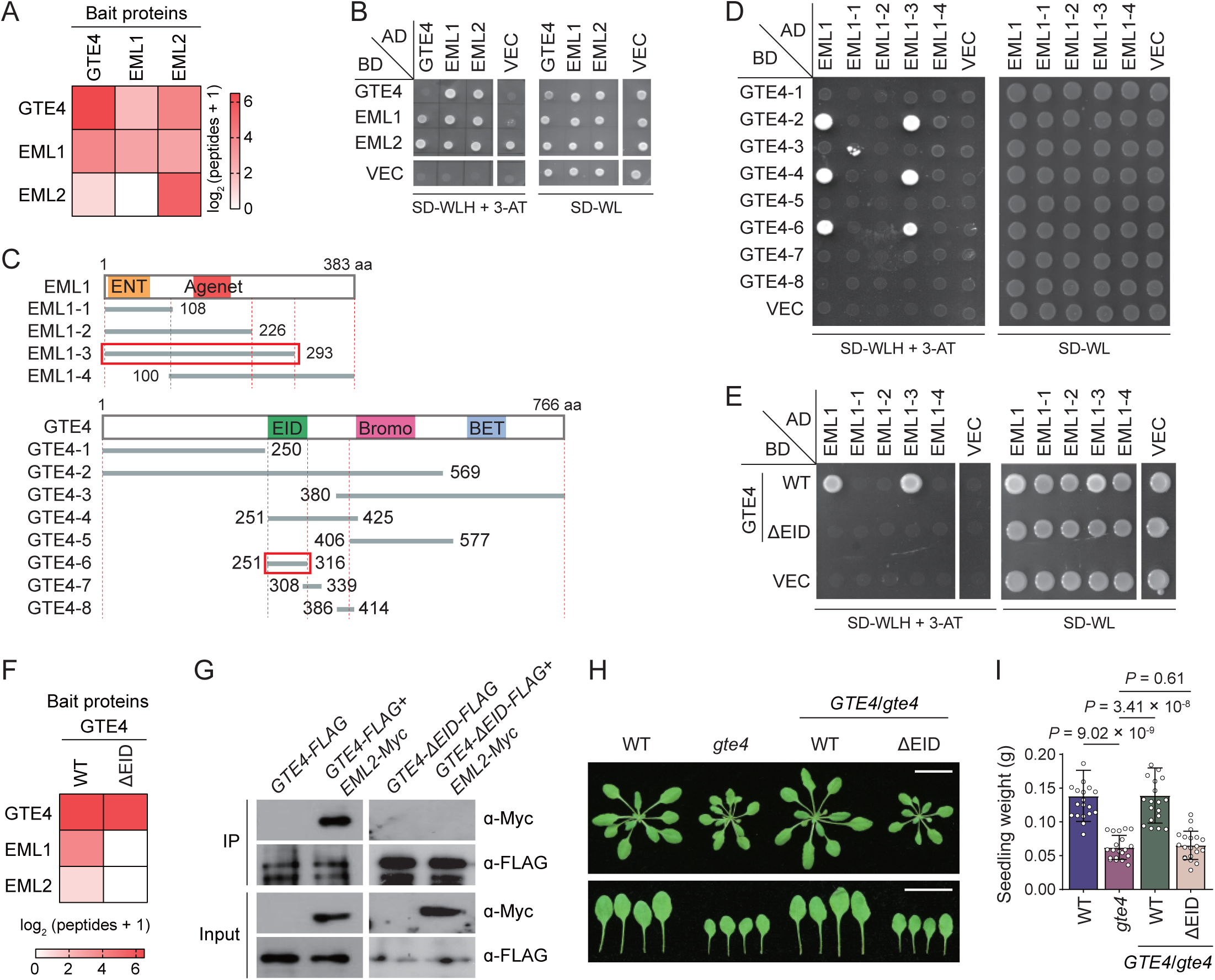
Determination of the interaction between GTE4 and EML1 or EML2. (A) Identification of EML1 and EML2 as GTE4-interacting proteins by AP-MS. The AP-MS experiment was performed using anti-FLAG antibody in *GTE4-FLAG*, *EML1-FLAG*, and *EML2-FLAG* transgenic plants. (B) Determination of the interaction between GTE4 and EML1 or EML2 by Y2H assays. The yeast strains expressing GAL4-AD and GAL4-BD-fused proteins were grown on SD-WLH supplemented by 3 mM 3-AT for determining the protein-protein interaction. The same yeast strains were grown on SD-WL as controls. (C) Schematic diagrams of full-length and truncated versions of GTE4 and EML1. The length and conserved domains of GTE4 and EML1 are indicated. Both the N- and C-terminal positions of each truncated version are labeled. The domains responsible for the GTE4-EML1 interaction as determined by Y2H were marked by red boxes. (D) Identification of the specific domains responsible for the GTE4-EML1 interaction by Y2H. A series of truncated versions of GTE4 and EML1 were used in the Y2H assay. The yeast strains expressing GAL4-AD and GAL4-BD-fused proteins were grown on SD-WLH supplemented by 3 mM 3-AT and on SD-WL. (E) Effect of the EID deletion on the interaction of GTE4 with EML1. The Y2H assay was performed to determine the interaction of full-length and EID-deleted GTE4 (ΔEID) with the full-length and truncated versions of EML1. (F) Determination of the effect of EID deletion on the interaction of GTE4 with EML1 and EML2 by AP-MS in Arabidopsis plants. Full-length and EID-deleted *GTE4-FLAG* transgenic plants were subjected to AP-MS using anti-FLAG antibody. (G) Determination of the effect of EID deletion on the GTE4-EML2 interaction by co-IP in Arabidopsis plants. Full-length and EID-deleted *GTE4-FLAG* transgenic plants were separately crossed to *EML2-Myc* transgenic plants, and the F1 plants harboring both the *FLAG*- and *Myc*-tagged transgenes were subjected to co-IP. (H,I) Effect of the EID deletion on the function of GTE4 in Arabidopsis plants. Full-length and EID-deleted *GTE4-FLAG* transgenes were transformed into the *gte4* mutant for complementation testing. The morphological phenotype of 21-day-old-plants, including the 3^rd^, 4^th^, 5^th^, and 6^th^ rosette leaves (H), and the statistical analysis of 21-day-old seedling weight (I) are shown. The values of plant weight of 21-day-old seedlings are shown as means ± SD (n = 20). P values were determined by Student t test with two tails. Bar, 2 cm.

To characterize the GTE4-EML complex, we determined the interaction between GTE4 and EML1/2 by yeast two-hybrid (Y2H) assays. The Y2H results indicated the interaction between GTE4 and EML1/2 (Figure 2B). Although the Y2H assay also identified the interaction between EML1 and EML2 (Figure 2B), the interaction between the EML proteins need to be validated in Arabidopsis plants. To dissect the domains responsible for the GTE4-EML1 interaction, we conducted Y2H assays using truncated versions of GTE4. These assays allowed us to identify a previously uncharacterized domain of GTE4 (amino acids 251-316) that forms a conserved α-helix (Supplemental Figure 4) as the region responsible for interacting with EML1 (Figure 2C, 2D). Importantly, the removal of this domain from GTE4 disrupted its interaction with EML1 (Figure 2E). Therefore, we named this domain the EML-interacting domain (EID). Additionally, the Y2H assays indicated that the N-terminal region of EML1 (amino acids 1-293) containing the ENT and Agenet domains is responsible for interacting with GTE4 (Figure 2C-2E).

To determine the impact of EID deletion on the interaction between GTE4 and EML1/2 in Arabidopsis plants, we generated transgenic plants expressing both the wild-type GTE4 and a version of GTE4 with the EID domain deleted (GTE4-ΔEID) (Supplemental Figure 2). We then performed AP-MS to determine whether EML1/2 could be co-purified with wild-type GTE4 or GTE4-ΔEID. The AP-MS result indicated that EML1/2 were co-purified with wild-type GTE4 but not with GTE4-ΔEID (Figure 2F; Dataset 2). The effect of EID deletion on the GTE4-EML2 interaction was further confirmed through co-immunoprecipitation assays (Figure 2G). These results indicate that the EID domain is responsible for the GTE4-EML1/2 interaction in Arabidopsis plants. To further validate the functional significance of the GTE4-EML1/2 interaction, we conducted complementation testing. We introduced the wild-type *GTE4* transgene as well as the *GTE4-ΔEID* transgene into the *gte4* mutant and assessed their ability to restore the developmental defects observed in the mutant. Remarkably, the wild-type *GTE4* transgene successfully rescued the developmental defects, while the *GTE4-ΔEID* transgene failed to do so (Figure 2H, 2I; Supplemental Figure 2, 5). These results suggest that the GTE4-EML1/2 interaction is required for the function of GTE4 in Arabidopsis plants.

### GTE4 and EML1/2 co-regulate development and gene expression

To investigate the functional association between EML1/2 and GTE4 in Arabidopsis plants, we examined the phenotypes of *eml1* and *eml2* single mutants. We found that the available *eml1* (SALK_077088) and *eml2* (SALK_116222) single mutants did not display any visible developmental defects compared to the wild type (data not shown), suggesting a potential redundancy between EML1 and EML2. We therefore generated an *eml1 eml2* (*eml1/2*) double mutant by genetic crossing and compared its developmental phenotypes with those of the *gte4* mutant. We found that the *eml1/2* double mutant exhibited similar developmental defects with the *gte4* mutant, including smaller rosette size, reduced seedling weight, and jagged leaves (Figure 3A-3C). To determine the genetic relationship between *gte4* and *eml1/2*, we generated a *gte4 eml1/2* (*gte4/eml1/2*) triple mutant by genetic crossing and then compared the phenotypes of *gte4*, *eml1/2*, and *gte4 eml1/2* mutants. We found that the *gte4/eml1/2* mutant exhibited more severe developmental defects compared to either *gte4* or *eml1/2* mutants alone (Figure 3A-3C). These results support the inference that GTE4 and EML1/2 have a synergistic effect on plant development.

**Figure 3.**
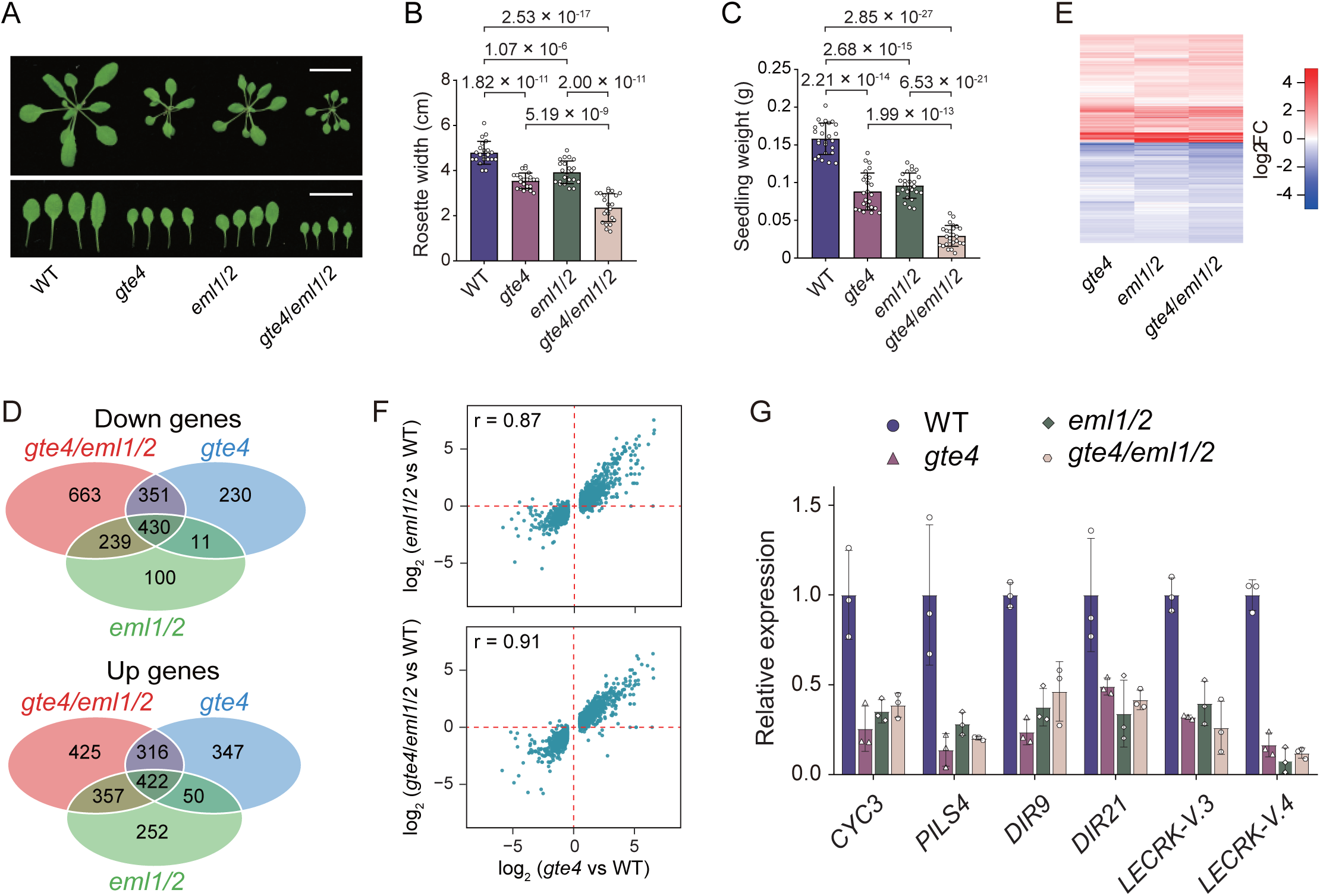
GTE4 and EML1/2 co-regulate plant development and gene expression. (A-C) Effect of *gte4*, *eml1/2*, and *gte4/eml1/2* mutations on plant development. The morphological phenotype, including the 3^rd^, 4^th^, 5^th^, and 6^th^ rosette leaves (A), rosette width (B), and seedling weight (C) are indicated in 21-day-old plants of wild-type, *gte4*, *eml1/2*, and *gte4/eml1/2* mutants. The values of rosette width and plant weight of 21-day-old seedlings are shown as means ± SD (n > 20). P values were determined by Student t test with two tails. Bar, 2 cm. (D) The overlaps of DEGs identified in *gte4*, *eml1/2*, and *gte4/eml1/2*, relative to the wild type. The overlaps of down-(upper panel) and up-regulated DEGs (lower panel) are independently shown. (E) Heatmaps showing the effect of *gte4*, *eml1/2*, and *gte4/eml1/2* on the expression of the combined set of DEGs identified in *gte4*, *eml1/2*, and *gte4/eml1/2*. The log_2_(fold change of reads between mutant and wild type) values are represented by color bars. (F) Scatter plots showing the correlation between the effects of *gte4*, *eml1/2*, and *gte4/eml1/2* on gene expression. The DEGs identified in the *gte4* mutant were subjected to the analysis. (G) Relative expression levels of representative genes as determined by RNA-seq. The gene expression levels in the mutants were normalized to those in the wild-type control Col-0. The values are shown as means ±SD (n = 3).

To explore how GTE4 and EML1/2 regulate development, we performed RNA deep sequencing (RNA-seq) to examine the impact of *gte4*, *eml1/2*, and *gte4/eml1/2* on gene expression. Analysis of the RNA-seq data revealed a significant number of differentially expressed genes (DEGs) in the three mutants compared to the wild type (log_2_FC > 0.5 or < -0.5, FDR < 0.05) (Figure 3D; Dataset 3). There was a substantial overlap of both up- and down-regulated genes among the three mutants (Figure 3D). Heat maps demonstrated that the majority of DEGs identified in the mutants were co-regulated (Figure 3E). Scatter plotting analysis revealed a high positive correlation in the expression levels of DEGs between *gte4* and *eml1/2* mutants, as well as between *gte4* and *gte4/eml1/2* mutants (Figure 3F). These results suggest that GTE4 and EML1/2 cooperate in the regulation of gene expression.

Among the co-regulated genes in *gte4*, *eml1/2*, and *gte4/eml1/2* mutants, we observed a significant reduction in the expression of *CYC3*, a cyclin family gene (Wang et al., 2004), and *PILS4*, an auxin transport-related gene (Barbez et al., 2012) (Figure 3G). This provides a potential explanation for the reduced size observed in these mutants. Interestingly, we also found a significant decrease in the expression of disease resistance-related genes, such as *DIR9*, *DIR21*, *LECRK-V.3*, and *LECRK-V.4* (Bouwmeester and Govers, 2009; Paniagua et al., 2017), in all three mutants (Figure 3G). This is consistent with previous studies showing the involvement of EML1 and EML2 in plant defense against pathogens (Tsuchiya and Eulgem, 2011; Coursey et al., 2018). Further investigations are needed to elucidate the precise mechanisms by which GTE4 and EML1/2 cooperate to regulate the expression of genes involved in specific biological processes.

### GTE4 and EML1/2 co-occupy genes enriched with histone acetylation and H3K4me3

To determine whether GTE4 and EML1/2 co-occupy chromatin to regulate gene transcription, we performed chromatin immunoprecipitation followed by deep sequencing (ChIP-seq) using *GTE4-FLAG* and *EML2-FLAG* transgenic plants. We obtained two independent replicates of the ChIP-seq data, which revealed 13,775 and 14,231 peaks for GTE4 and EML2, respectively (Dataset 4 and 5). We found that the majority of both GTE4 and EML2 peaks were found in intragenic regions near the transcription start site (TSS) of protein-coding genes (Figure 4A-4C). By analyzing the chromatin annotations of these peaks, we identified 13,371 genes bound by GTE4 and 13,797 genes bound by EML2. There was a significant overlap between the genes bound by GTE4 and EML2 (Figure 4D), indicating a co-occupancy of GTE4 and EML1/2 at TSS-proximal intragenic regions at the whole-genome level.

**Figure 4.**
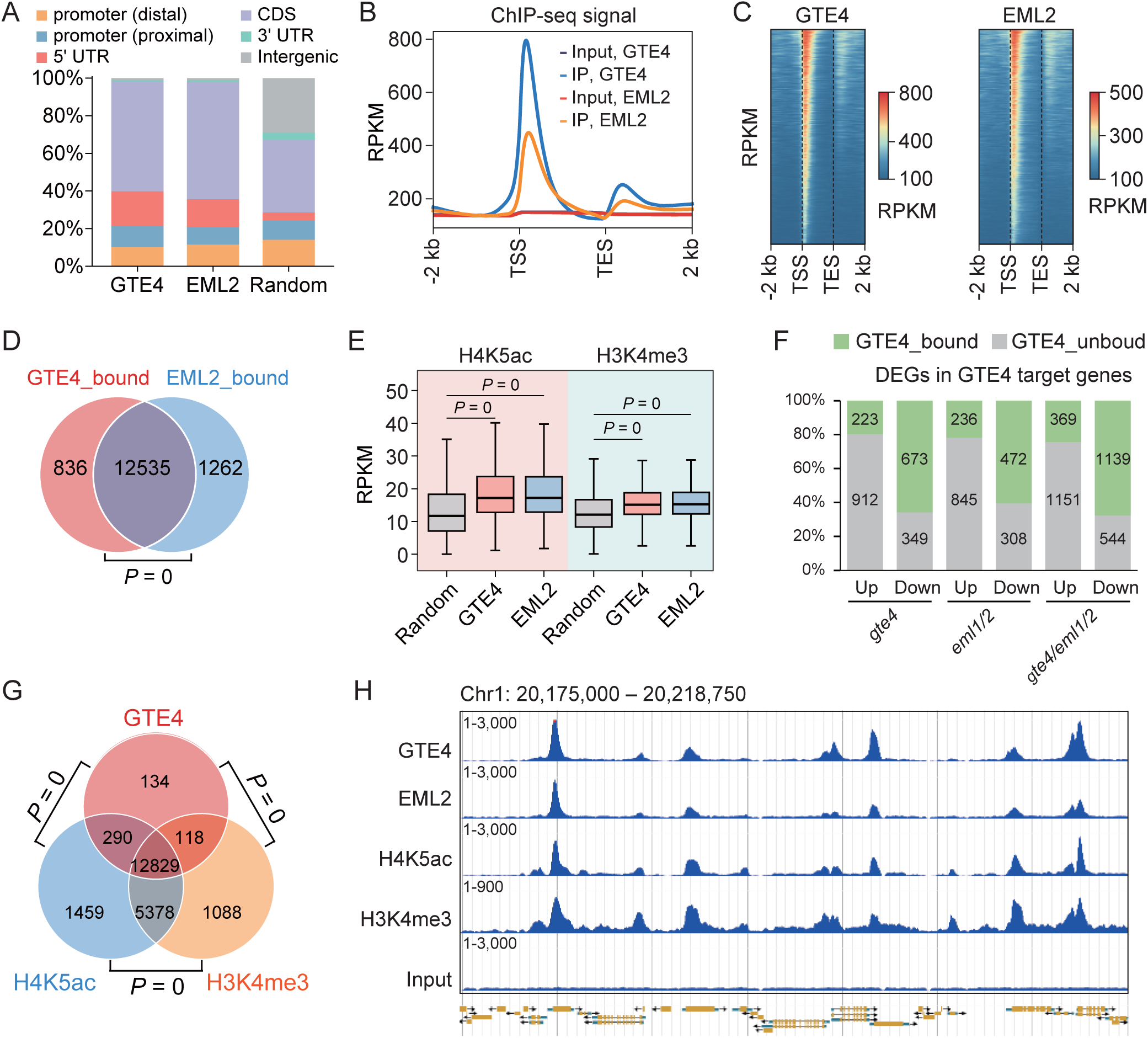
GTE4 and EML2 co-occupy target genes enriched with both H4K5ac and H3K4me3. (A) Distribution of GTE4 and EML2 ChIP-seq peaks at different genomic regions. Proximal and distal promoter regions represent 1-400 bp and 401-1000 bp regions upstream to the transcription start site, respectively. (B,C) Distribution of GTE4 and EML2 ChIP-seq signals over the GTE4- and EML2-enriched genes. Metaplots (B) and heatmaps (C) are independently shown. In heatmaps, the GTE4- and EML2-enriched genes were sorted by the enrichment levels of ChIP-seq signals. RPKM, reads per kilobase per million mapped reads; TSS, transcription start site; TES, transcription end site. (D) Venn diagram showing the overlap between GTE4- and EML2-bound genes. P value determined by one-tailed hypergeometric test indicates the significance of overlap. (E) The H4K5ac and H3K4me3 ChIP-seq levels of GTE4-bound genes, EML2-bound genes, and random genes in the wild-type plants. P values were determined by two-tailed, Mann Whitney U test (unpaired). (F) The ratio of GTE4-bound DEGs to total DEGs identified in *gte4*, *eml1/2*, and *gte4/eml1/2* mutants. The up- and down-regulated DEGs were independently analyzed. (G) The overlaps among GTE4-enriched genes, H4K5ac-enriched genes, and H3K4me3-enriched genes. The significance of overlapping was indicated by P values as determined by one-tailed hypergeometric test. (H) Genome viewers showing the ChIP-seq signals for GTE4, EML2, H4K5ac, and H3K4me3 at representative genomic region. The scales of RPKM are shown for ChIP-seq signals.

Given that GTE4 binds to acetylated histone and EML1/2 binds to H3K4me3, we assessed whether GTE4- and EML2-bound genes have high levels of H4ac and H3K4me3. By analyzing previously published H4K5ac and H3K4me3 ChIP-seq data (Shang et al., 2021; Zheng et al., 2023), we found that GTE4- and EML2-bound genes exhibited significantly higher levels of H4ac and H3K4me3 than randomly selected genes (Figure 4E; Dataset 6). The enrichment levels of GTE4 were increased in gene deciles with incremental levels of H4K5ac and H3K4me3, except for the last decile (Supplemental Figure 6A, 6B). These findings suggest that GTE4 binds to acetylated histone and EML1/2 binds to H3K4me3, contributing to their association with chromatin.

By comparing RNA-seq and ChIP-seq data, we found that GTE4- and EML2-bound genes were highly enriched at down-regulated genes but not at up-regulated genes, as identified in the *gte4*, *eml1/2*, and *gte4/eml1/2* mutants (Figure 4F). Although a limited number of genes enriched solely with either H4K5ac or H3K4me3 showed occupied by GTE4, the majority of genes enriched with both H4K5ac and H3K4me3 were indeed occupied by GTE4 (Figure 4G, 4H; Supplemental Figure 6C). These results strongly suggest that GTE4 and EML1/2 have a preference for binding to genes enriched with both histone acetylation and H3K4me3, ultimately facilitating the transcription of these specific genes.

### The binding of GTE4 and EML1/2 to chromatin is partially interdependent

To investigate the impact of EML1/2 on the chromatin binding of GTE4, we conducted ChIP-seq using *GTE4-FLAG* transgenic plants in the *eml1/2* mutant and the wild type (Supplemental Figure 7A). Our analysis indicated that although the GTE4-bound genes identified in the *eml1/2* mutant and the wild type displayed similar abundance and had a significant overlap (Figure 5A; Dataset 4), the enrichment of GTE4 at its target genes was significantly lower in the *eml1/2* mutant than in the wild type (Figure 5B-5E; Dataset 7). These findings suggest that EML1/2 are involved in the association of GTE4 with chromatin.

**Figure 5.**
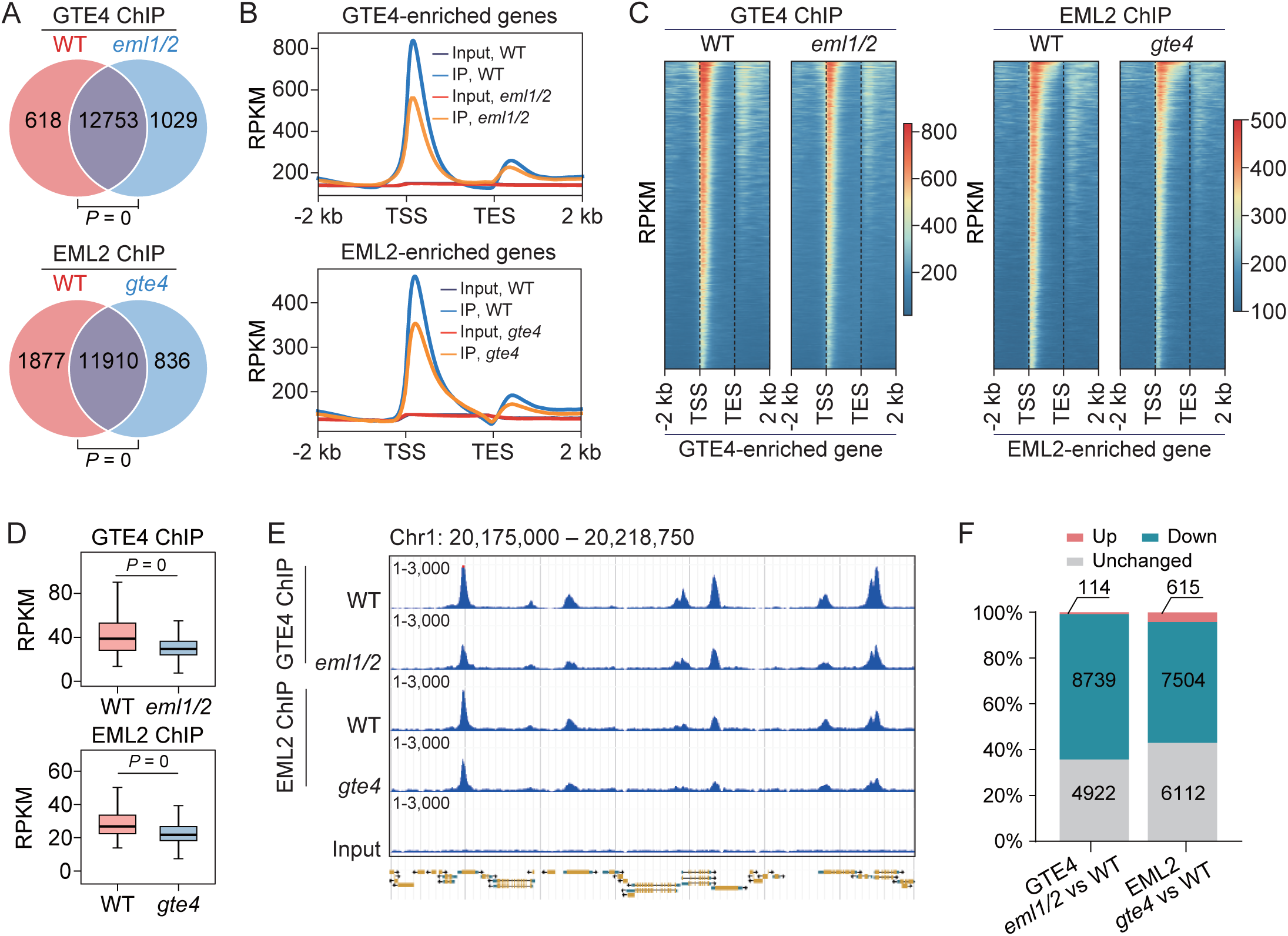
GTE4 and EML1/2 exhibit a partial interdependent binding to their target genes. (A) Venn diagrams showing the overlap of GTE4-bounds genes between wild type and *eml1/2* mutant (upper panel) and the overlap of EML2-bound genes between wild type and *gte4* mutant (lower panel). P values were determined by one-tailed hypergeometric test. (B) Determination of the effect of *eml1/2* on GTE4 ChIP-seq signals and the effect of *gte4* on EML2 ChIP-seq signals. Meta plots show the average GTE4 and EML2 ChIP-seq signals in the wild type, *gte4*, or *eml1/2* mutants. (C) Heat maps showing the distribution patterns of GTE4 ChIP-seq signals in the wild type and *eml1/2* mutant and the distribution of EML2 ChIP-seq signals in the wild type and *gte4* mutant. The genes were sorted based on the degree of enrichment of GTE4 and EML2 on their respective target genes. (D) Box plots showing the effect of *eml1/2* on the enrichment of GTE4 and the effect of *gte4* on the enrichment of EML2. The enrichment levels of GTE4 and EML2 were determined at GTE4 and EML2 ChIP-seq peaks, respectively. P values were determined by two-tailed, paired Mann Whitney U test. (E) Genome viewers showing the effect of *eml1/2* on GTE4 enrichment and the effect of *gte4* on EML2 enrichment at the representative genomic region. The scale of RPKM is shown. (F) Bar charts showing the quantities of GTE4-bound genes with *eml1/2*-affected enrichment and the quantities of EML2-bound genes with *gte4*-affected enrichment. GTE4-bound genes identified in either the wild type or *eml1/2* mutant were combined for determining the effect of *eml1/2* on the enrichment of GTE4; EML2-bound genes identified in either the wild type or *gte4* mutant were combined for determining the effect of *gte4* on the enrichment of EML2.

To further explore the relationship between GTE4 and EML2, we performed ChIP-seq using *EML2-FLAG* transgenic plants in the *gte4* mutant and the wild type (Supplemental Figure 7B). Similarly, we found that the EML2-bound genes identified in the *gte4* mutant exhibited comparable abundance to those in the wild type and displayed a high degree of overlap (Figure 5A; Dataset 5). However, the enrichment of EML2 at its target genes was significantly reduced in the *gte4* mutant compared to the wild type (Figure 5B-5E; Dataset 7), suggesting that GTE4 is involved in the association of EML1/2 with chromatin.

Although the *eml1/2* and *gte4* mutants showed diminished chromatin binding of GTE4 and EML2, respectively, the binding was not completely abolished (Figure 5B-5E). This suggests that the binding of GTE4 and EML1/2 to chromatin is partially dependent on each other. Furthermore, we observed that the majority of GTE4 and EML2 target genes identified in the wild type exhibited reduced enrichment in the *eml1/2* and *gte4* mutants, respectively, whereas the enrichment of GTE4 and EML2 was not significantly affected in the mutants at a subset of their target genes (Figure 5E, 5F). These findings confirm that the binding of GTE4 and EML1/2 to chromatin is partially interdependent.

### The binding of GTE4 to H4ac is involved in GTE4 binding to chromatin

Although the binding of the GTE4 bromodomain to acetylated histone has been demonstrated (Figure 1B-1D), it remains unclear whether this binding is necessary for the binding of GTE4 to chromatin *in vivo*. To investigate the impact of the Y497A/N498A mutation on the binding of GTE4 to chromatin, we performed ChIP-seq using transgenic plants expressing wild-type GTE4 and the mutated form GTE4-Y497A/N498A (Supplemental Figure 2). By comparing the ChIP-seq data for the wild type and the mutated GTE4, we observed a significant overlap in the genes bound by both forms of GTE4 (Figure 6A; Dataset 8). However, at the wild type GTE4-bound chromatin regions, the overall enrichment level of GTE4-Y497A/N498A was substantially lower than that of the wild-type GTE4 (Figure 6B-6E; Dataset 9), indicating that the conserved bromodomain is required for the association of GTE4 with chromatin at the whole-genome level. Moreover, we found that the Y497A/N498A mutation led to a significant reduction of GTE4 enrichment at most of GTE4-bound genes (Figure 6F, 6G), supporting the idea that the bromodomain binding to acetylated histone is required for the association of GTE4 with chromatin.

**Figure 6.**
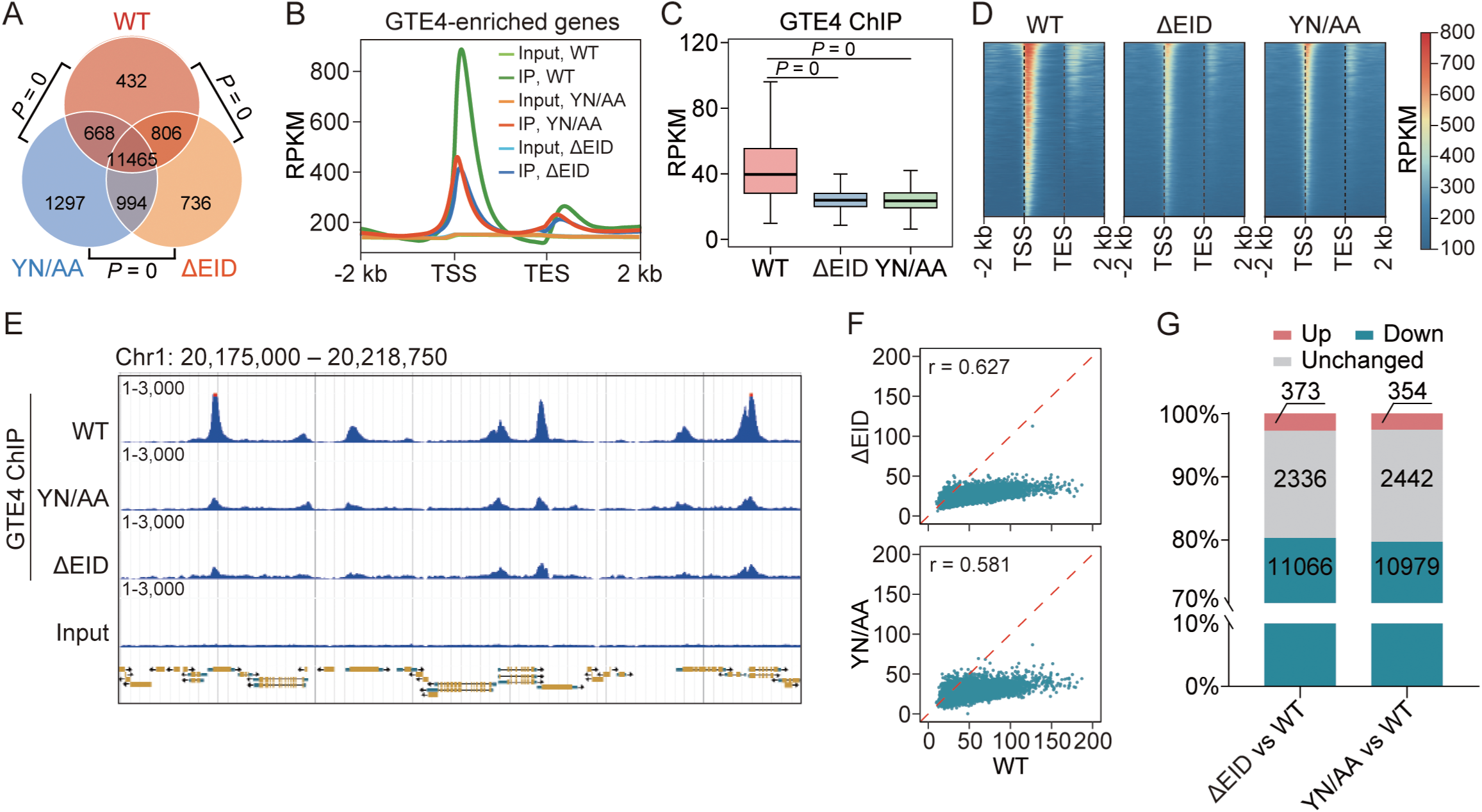
Both the bromodomain and EML-interacting domain are involved in the binding of GTE4 to chromatin. (A) The overlaps of genes bound by wild-type GTE4, GTE4-Y497A/N498A, and GTE4-ΔEID. P values were determined by one-tailed hypergeometric test. (B-D) The effects of Y497A/N498A mutation and EID deletion on the enrichment of GTE4 at its target genes. The effects were indicated by meta plots (B), box plots (C), and heat maps (D). The meta plots show the average ChIP-seq signals over the wild-type GTE4-bound genes. Box plots indicate the enrichments of wild-type GTE4, GTE4-Y497A/N498A, and GTE4-ΔEID at the wild-type GTE4 ChIP-seq peaks. P values shown in the box-plot chart were determined by two-tailed, paired Mann Whitney U test. Heat maps show the distribution of ChIP-seq signals for wild-type GTE4, GTE4-Y497A/N498A, and GTE4-ΔEID over GTE4-bound genes. The genes were sorted by the degree of GTE4 enrichment at its target genes. TSS, transcription start site; TES, transcription end site. (E) Genome viewers showing the effects of Y497A/N498A mutation and EID deletion on GTE4 enrichment at the representative genomic region. The scale of RPKM is shown. (F) Scatter plots showing the comparison between the enrichment of wild-type GTE4 and the enrichments of GTE4-Y497A/N498A or GTE4-ΔEID at their target genes. (G) Bar charts showing the numbers of genes with up- and down-regulated levels of GTE4 enrichment caused by the Y497A/N498A mutation and the EID deletion.

Given that EML1 has been shown to bind to H3K4me3 (Zhao et al., 2018), we aimed to investigate whether the association of GTE4 with chromatin is affected when the GTE4-EML1/2 interaction is disrupted by deleting the EID domain. To assess the impact of the EID deletion on the association of GTE4 with chromatin, we performed ChIP-seq for GTE4-ΔEID using transgenic plants expressing GTE4-ΔEID-FLAG (Supplemental Figure 2). Our analysis revealed that the genes bound by GTE4-ΔEID significantly overlapped with those bound by wild-type GTE4 (Figure 6A; Dataset 8). However, at the loci where wild-type GTE4 was bound, the overall enrichment of GTE4-ΔEID was notably reduced compared to wild-type GTE4 (Figure 6B-6E; Dataset 9), indicating that the EID domain is necessary for the association of GTE4 with chromatin. Moreover, we observed a significant decrease in GTE4 enrichment at most of its bound loci when the EID domain was deleted (Figure 6F, 6G). Interestingly, the effect of the EID deletion on GTE4 enrichment was stronger than that of the *eml1/2* mutation (Figure 5B-5F, 6B-6G), suggesting that the EID domain also participates in the association of GTE4 with chromatin in an EML1/2-independent manner. Considering that there are two additional EML homologs (EML3 and EML4) in Arabidopsis, it is possible that EML3 and/or EML4 may function redundantly with EML1/2 to enhance the association of GTE4 with chromatin.

## Discussion

The tandem PHD-Bromodomain module has been found in numerous chromatin-related proteins in yeast and metazoans (Ito et al., 1999; Schultz et al., 2001; Ruthenburg et al., 2007; Plotnikov et al., 2014; Tallant et al., 2015; Bardhan et al., 2023). BPTF, a core subunit of the human NURF chromatin-remodeling complex, contains a PHD-Bromodomain module that facilitates coordinated recognition of histone acetylation and H3K4me3 (Ruthenburg et al., 2011). Although a PHD-Bromodomain module was reported in MBD9, a component of SWR1 complex responsible for H2A.Z deposition, in Arabidopsis (Nie et al., 2019; Potok et al., 2019; Sijacic et al., 2019; Luo et al., 2020), the disruption of PHD or bromodomain did not affect the biological function of MBD9 (Luo et al., 2020). It remains unclear whether and how the coordination between histone acetylation and H3K4me3 plays a crucial role in plants. In this study, we found that the acetylated histone-binding protein GTE4 forms a complex with the H3K4me3-binding EML1/2, revealing a previously uncharacterized coordination between the two histone modifications. Considering the well-established roles of histone acetylation and H3K4me3 in transcriptional activation (Struhl, 1998; Bannister and Kouzarides, 2011), the combinatorial recognition of the two histone modifications likely synergistically contributes to the activation of transcription. GTE4 was previously thought to have both activating and repressive roles in transcriptional regulation (Zhou et al., 2022). However, through the analysis of RNA-seq and ChIP-seq data, we discovered that both GTE4 and EML occupy genes that are predominantly down-regulated DEGs in the *gte4* and *eml1/2* mutants, supporting the notion that GTE4 primarily functions in transcriptional activation.

We found that the bromodomain of GTE4 exhibits a strong binding affinity for H4 with multiple acetylated lysine sites. Previous studies have investigated the binding specificity of bromodomains to different acetylated lysine sites on histones in various BET proteins of other eukaryotes (Owen et al., 2000; Zeng and Zhou, 2002). Structural analyses have revealed how a mono bromodomain can simultaneously bind to polyacetylated histone lysine sites (Moriniere et al., 2009). In our current study, we observed that the single bromodomain of GTE4 has significantly higher binding affinity for polyacetylated histone peptides compared to their monoacetylated counterparts. This suggests that the recognition of polyacetylated histone lysine sites by a mono bromodomain is conserved in plants. The strong binding affinity of GTE4 for polyacetylated H4 likely contributes to its association with chromatin exhibiting high levels of histone acetylation. Previously, EML1 was identified as a H3K4me3-binding protein, and the Agenet/Tudor domain of EML1 was found to act as a plant-specific H3K4me3-binding module (Zhao et al., 2018). Our study suggests that the GTE4-EML complex co-recognizes histone acetylation and H3K4me3 on their shared target genes, thereby establishing a connection between the two histone modifications in plants.

Although our study indicates that GTE4 and EML1/2 form a complex in Arabidopsis, we cannot exclude the possibility that either the GTE4 homologs or the EML1/2 homologs are also involved in the formation of redundant GTE-EML complexes. In addition to GTE4, several GTE4 homologs, including GTE3, GTE7, GTE8, GTE9, GTE10, and GTE11, were co-purified with EML2, as determined by AP-MS (Dataset 2). Similarly, in addition to EML1/2, the EML1/2 homolog EML3 was co-purified with GTE4 (Dataset 2). These results support the inference that the GTE4 and EML1/2 homologs form redundant GTE-EML complexes. The functional redundancy of the GTE4 and EML1/2 homologs provides a plausible explanation for the finding that only a small part of GTE4- and EML1/2-occupied genes are differentially expressed in the *gte4* and *eml1/2* mutants. It is also possible that the differentially expressed genes in the *gte4* and *eml1/2* mutants can only be identified during specific developmental stages or under certain adverse environmental conditions but not under the conditions tested in the current study.

Previous studies indicated that GTE4 and EML proteins are involved in the regulation of plant development (Milutinovic et al., 2019) and disease resistance (Tsuchiya and Eulgem, 2011; Coursey et al., 2018; Zhou et al., 2022), suggesting the GTE4-EML complex has multiple functions in Arabidopsis plants. We predict that the GTE-EML complexes are responsible for establishing a priming state for transcriptional activation rather than for directly mediating transcriptional activation, and certain uncharacterized transcription regulators are specifically responsible for the activation of different sets of GTE4- and EML1/2-occupied genes. Further studies are required to identify which transcription factors are involved in the activation of different sets of GTE4- and EML1/2-occupied genes and to determine how the transcription factors are coordinated with the GTE4-EML1/2 complex to regulate the activation of these genes.

## Materials and methods

### Plant materials and constructs

Arabidopsis plants used in this study are in the Columbia (Col-0) background. The T-DNA insertion mutants *gte4* (SALK_083697), *eml1* (SALK_077088), and *eml2* (SALK_116222) were obtained from Arabidopsis Biological Resource Center (ABRC). The *eml1/2* double mutant and the *gte4/eml1/2* triple mutant were generated by crossing. Arabidopsis seedlings were grown on Murashige and Skoog (MS) medium plates with a 16-h light and 8-h dark photoperiod at 22°C. Twelve-day-old plants were transferred to soil and grown in the culture room under the same photoperiod and temperature conditions.

The full-length *GTE4*, *EML1*, and *EML2* genomic DNA sequences driven by their own promoters were fused to *3×FLAG* tag or 3*×Myc* tags at their 3’-terminal regions and cloned into the modified pCAMBIA1305 vector. The full-length and *EID*-deleted versions of *GTE4* coding sequences driven by the native *GTE4* promoter was also fused to *3×FLAG* tag at their 3’-terminal regions and cloned into the modified pCAMBIA1305 vector. The primers used for generating the constructs are shown in Dataset 10. The constructs were transformed into the *Agrobacterium* strain GV3101 and introduced into Arabidopsis plants via the floral-dip method. The T1 transgenic seedlings were selected on MS medium supplemented by 30 mg/L hygromycin and 50 mg/L ampicillin. The transgenic plants were subjected to affinity purification followed by mass spectrometry. The *EID*-deleted *GTE4* fragment was amplified from the *pGTE4::GTE4-FLAG* construct and cloned into the pCAMBIA1305 vector, thereby obtaining the *pGTE4::GTE4-ΔEID-FLAG* construct. The *Y497A*/*N498A* mutations were introduced into *GTE4* using the Mut Express MultiS Fast Mutagenesis Kit (Vazyme, C215-01).

### Yeast two-hybrid assay

The full-length and a series of truncated sequences of *GTE4*, *EML1*, and *EML2* cDNA were cloned into pGADT7 and pGBKT7 vectors. The pGADT7 and pGBKT7 constructs were transformed into the yeast strains AH109 and Y187, respectively. The strains AH109 grew on synthetic dropout medium lacking Leu (SD-L); the strain Y187 grew on synthetic dropout medium lacking Trp (SD-W). Then, the positive AH109 and Y187 strains were subjected to mating in YPDA (yeast peptone dextrose adenine) liquid medium for 16-20 hours. After mating, the culture was spread on a synthetic dropout medium plate lacking both Trp and Leu (SD-WL). The positive colonies grown on SD-WL medium were suspended and spotted on synthetic dropout medium lacking Trp, Leu, and His (SD-WLH) supplemented by 3 mM 3-amino-1,2,4-triazole (3-AT) and on SD-WL.

### Affinity purification, mass spectrometry, and co-immunoprecipitation

Three grams of plant material grown on MS medium was harvested and ground into fine powder in liquid nitrogen. The powder was suspended in 10 ml lysis buffer (50 mM Tris-HCl [pH 7.5], 150 mM NaCl, 5 mM MgCl_2_, 10% glycerol, 0.1% NP-40, 0.5 mM DTT, 1 mM PMSF, and 1% Roche protease inhibitor cocktail) and rotated for 30 min at 4℃, followed by centrifugation at 12,000 *g* for 10 min. The supernatant containing total plant proteins was passed through Miracloth (Millipore, 475855-1R) and incubated with 100 μl of anti-FLAG M1 Agarose affinity gel (Sigma, A4596) for 2 hours. The beads were harvested by centrifugation at 150 *g* for 3 min at 4°C and then washed four times with lysis buffer. The proteins bound to the beads were eluted with 3×FLAG peptide (Sigma, F4799) for 30 min at 4 °C. The eluate was run on a 10% SDS-PAGE gel, followed by silver staining with the ProteoSilver Silver Stain Kit (Sigma, PROT-SIL1).

Acetylated-histone-binding proteins were identified from Arabidopsis plants as previously described (Qian et al., 2021). In brief, total proteins were extracted from 2 grams of wild-type plants and then incubated with 2 μg of biotin-labeled H4K5/8/12/16ac (1-20 aa) peptides (Scilight-Peptide, Beijing) for 3 hours at 4℃ with rotation. 40 μl of Streptavidin MagneSphere Paramagnetic Particles (Promega, Z5481) were added to the mixture and incubated for 1 hour at 4℃ with rotation. After the particles were washed three times with wash buffer 1 (320 mM NaCl, 50 mM Tris-HCl [pH 8.0], and 0.5% NP-40) and three times with wash buffer 2 (150 mM NaCl, and 50 mM Tris-HCl [pH 8.0]), they were suspended and boiled in 1×SDS sample buffer and run on a 10% SDS-PAGE gel, followed by silver staining.

The mass spectrometry analysis was performed as previously described (Tan et al., 2020). In brief, the silver-stained protein bands were excised, de-stained, and digested with trypsin at 37℃ overnight. The peptides were eluted on a capillary column and sprayed into a linear trap quadrupole mass spectrometer equipped with a nano-ESI ion source (Thermo Fisher Scientific). The obtained spetra were searched on the Mascot server (Matrix Science Ltd., London, UK) against the IPI (International Protein Index) Arabidopsis protein database.

For co-IP, the total proteins extracted from Arabidopsis plants were subjected to immunoprecipitation using anti-FLAG M1 Agarose affinity gel (Sigma, A4596), followed by western blotting. *pEML2::EML2-Myc* transgenic plants were crossed to *pGTE4::GTE4-FLAG* and *pGTE4::GTE4-ΔEID-FLAG* transgenic plants, and the F1 plants harboring both the *Myc*- and *FLAG*-tagged transgenes were used for co-IP.

### RNA transcript analysis

Twenty-day-old wild type, *gte4*, *eml1/2*, and *gte4/eml1/2* plants grown in soil were used for RNA-seq analysis. One gram of plant material was subjected to total RNA extraction using Trizol reagent (Invitrogen) strictly adhering to the manufacturer’s instructions. Subsequently, the extracted total RNA was sent to Berry Genomics Co., Ltd. for reverse transcription and library construction. The constructed libraries were then sequenced using an Illumina NovaSeq6000 instrument with a paired-end scheme (2 X 150 bp). After adapters and poor-quality reads were removed, the clean reads were mapped to the Arabidopsis genome (TAIR 10) with HISAT2 (v2.2.0) (Kim et al., 2019). The differentially expressed genes (DEGs) were identified with a filter (FDR < 0.05, |log_2_FC| > 0.5) with the R (v4.0.3) package edgeR (v3.32.1) (Robinson et al., 2010). The RNA-seq results shown in this study were from three independent biological replicates. The heatmap of DEGs was draw by using the R package gplots (v3.1.1). Venn diagrams were drawn by using the R package ggvenn (v0.1.9). Scatter plots were drawn using the R package ggplot2 (v3.3.5).

### Chromatin immunoprecipitation

Transgenic Arabidopsis plants used for ChIP-seq comprise: wild-type *GTE4-FLAG*, *GET4-Y497A/N498A*, and *GTE4-ΔEID-FLAG* transgenic plants in the *gte4* background; *GTE4-FLAG* transgenic plants in both wild-type and *eml1/2* backgrounds; and *EML2-FLAG* transgenic plants in both wild-type and *gte4* backgrounds. Four grams of 12-day-old seedlings were harvested and fixed with 1% formaldehyde (Sigma, F8775). Samples were ground in liquid nitrogen and homogenized in 40 ml of NEB1 buffer (10 mM Tris-HCl [pH 8.0], 0.4 M sucrose, 10 mM MgCl_2_, 5 mM DTT, 0.1 mM PMSF, 1% Roche protease inhibitor cocktail tablet). The homogenate was filtrated through two layers of Miracloth, and then centrifuged at 2,000 *g* for 20 min at 4°C. After washing four times with NEB2 buffer (10 mM Tris-HCl [pH 8.0], 0.25 M sucrose, 10 mM MgCl_2_, 1% Triton X-100, 0.1 mM PMSF), the nuclei were resuspended in 1.2 ml of NEB3 buffer (10 mM Tris-HCl [pH 8.0], 1.7 M sucrose, 2 mM MgCl_2_, 0.15% Triton X-100, 0.1 mM PMSF, 1% Roche protease inhibitor cocktail tablet). Subsequently, 600 μl of NEB3 was added to a new tube and overlaid with the sample solution before centrifugation at 14,000 *g* for 1 hour at 4°C. The pellet was resuspended in 400 μl of nuclear lysis buffer (50 mM Tris-HCl [pH 8.0], 10 mM EDTA, 1% SDS, 0.1 mM PMSF, 1% Roche protease inhibitor cocktail tablet), and diluted by adding 800 μl of dilution buffer (16.7 mM Tris-HCl [pH 8.0], 1.2 mM EDTA, 167 mM NaCl, 1.1% Triton X-100, 0.1 mM PMSF, 1% Roche protease inhibitor cocktail tablet). The sample was then sonicated using Bioruptor (Diagenode). Following centrifugation at 16,000 *g* for 15 min at 4°C, the supernatant was diluted by adding 2.8 ml of dilution buffer. After preclearing using protein G beads (Thermo Fisher Scientific), the chromatin was immunoprecipitated using anti-FLAG antibody (SIGMA, F1804) and then incubated with protein G beads. The slurry was washed twice with low salt buffer (20mM Tris-HCl [pH 8.0], 2 mM EDTA, 150 mM NaCl, 0.1% SDS, 1% Triton X-100, 0.1 mM PMSF), high salt buffer (20mM Tris-HCl [pH 8.0], 2 mM EDTA, 500 mM NaCl, 0.1% SDS, 1% Triton X-100, 0.1 mM PMSF), LiCl wash buffer (10mM Tris-HCl [pH 8.0], 1 mM EDTA, 0.25 M LiCl, 1% sodium deoxycholate, 1% NP-40, 0.1 mM PMSF), and TE buffer (10mM Tris-HCl [pH 8.0], 1 mM EDTA, 0.1 mM PMSF). The protein-DNA complex was eluted with elution buffer (0.1% SDS, 0.1 M NaHCO_3_), reverse cross-linked at 65°C, and treated with proteinase K and RNase. The DNA was purified and used to prepare DNA libraries with the NEXTflex™ Rapid DNA-Seq Kit (Bioo Scientific).

The libraries were sent to Berry Genomics Co., Ltd. and sequenced on an Illumina NovaSeq6000 instrument using a paired-end scheme (2 X 150 bp). After adapters and low-quality reads were removed, the clean reads were mapped to the Arabidopsis genome by Bowtie2 (v2.3.4) allowing one mismatch (Langmead and Salzberg, 2012). Probable PCR duplicates were removed by using Picard Tools (v2.23.0) with MarkDuplicates. For FLAG ChIP-seq, enriched peaks were identified by MACS2 (v2.2.7.1) using the input reads as a negative control (Zhang et al., 2008). The read counts were normalized to RPKM (reads per kilobase per million mapped reads) by the number of clean reads mapped to the genome in each library. The ChIP-seq results shown in this study are from two independent biological replicates. The boxplots and scatterplots were drawn by using the R package ggplot2 (v3.3.5). The profile plot and heatmap of ChIP-seq were drawn by using DeepTools (v3.5.1). Venn diagrams were drawn by using the R package ggvenn (v0.1.9).

### Protein purification, histone peptide array, and pull-down assay

The DNA sequences encoding the bromodomain of GTE4 and the full-length EML1 were cloned into the pGEX6p-1 vector in fusion with the 5’-terminal GST tag for protein expression. Site-directed mutagenesis was performed using Mut Express II fast mutagenesis Kit V2 (Vazyme Biotech) to express the mutated bromodomain versions of GTE4, in which the mutations include L452D, Y455A, A494D, and Y497A/N498A. The plasmids were transformed into the *E. coli* strain BL21 (Rosetta DE3). The positive strain was incubated in LB medium at 37℃ to OD 600 = 0.6, followed by incubation in LB medium supplemented with 0.1 mM IPTG overnight at 16°C. The bacterial cells were harvested by centrifugation and suspended by GST-lysis buffer (20 mM Tris-HCl [pH 7.5], 150 mM NaCl). The suspension was sonicated for 3 min and then subjected to centrifugation at 18,000 *g* for 1 hour at 4℃. The supernatant was incubated with Glutathione-Sepharose 4B (GE Healthcare, 71024800-EG). After the beads were washed with GST-lysis buffer, the GST-fused proteins bound to the beads were eluted using GST elution buffer (50 mM Tris-HCl [pH 7.5], 150 mM NaCl, and 20 mM glutathione).

The MODified Histone Peptide Array (Active Motif) was used to identify the histone marks recognized by EML1 as previously described (Zhang et al., 2016). The histone peptide array was blocked in TTBS buffer (10 mM Tris-HCl [pH 7.5], 150 mM NaCl and 0.05% Tween-20) containing 5% non-fat dried milk at room temperature (RT) for 1 hour, followed by three-time washing in TTBS buffer at RT. Subsequently, 50 μg of the GST-fused EML1 protein was incubated with the array in 10 ml of binding buffer (50 mM HEPES [pH 7.5], 50 mM NaCl, 5% glycerol, 0.4% BSA, 2 mM DTT) overnight at 4°C. The array was then washed three times in 20 ml of TTBS buffer, followed by incubation with anti-GST antibody in TTBS buffer for 2 hours at RT. After the array was washed five times with TTBS, it was subjected to immunoblotting and then analyzed using the Active Motif Array Analyze software.

For histone peptide pull-down assays, 2 μg of GST-fused protein was incubated with 1 μg of biotin-labeled histone peptides in 600 μl of binding buffer (50 mM Tris-HCl [pH 7.5], 150 mM NaCl, and 0.05% NP-40) for 4 hours at 4°C with rotation. 25 μl of Streptavidin MagneSphere Paramagnetic Particles (Promega, Z5481) was added to each tube and incubated for 2 hours at 4°C to capture histone peptides. A tube without addition of histone peptides served as a negative control. After the particles were washed with 1 ml binding buffer for 4 times, the proteins bound to the particles were dissolved in 1 X SDS sample buffer by boiling and then run on SDS-PAGE gel for immunoblotting. The GST-fused proteins without addition of histone peptides were used as negative controls. The histone peptides, including H3 (1-19 aa), H3K4/9/14/18ac (1-26 aa), and H3K14ac (1-19 aa), were commercially produced (Scilight-Peptide, Beijing). Additional histone peptides used in this study include H3 (1-21 aa) (Millipore, 12-403), H4 (2-24 aa) (Millipore, 12-372), H4K5/8/12/16ac (1-18 aa) (Millipore, 12-379), H3K36ac (EpiCypher, 12-0021), and H3K4me3 (Millipore, 12-564).

### ITC assay

The binding affinity of the bromodomain of GTE4 with histone peptides was assessed by a standard ITC assay. ITC binding curves were measured using a Microcal PEAQ-ITC instrument (Malvern). Purified proteins were dialyzed against ITC buffer containing 100 mM NaCl, 20 mM HEPES [pH 7.0], and 2 mM β-mercaptoethanol. The peptides were dissolved in the same buffer. Proteins were diluted to 0.04 mM with the ITC buffer, and 320 μl of the sample was added to the ITC instrument. Peptides were diluted to 0.6 mM and 60 ul of the peptides were added to the ITC instrument. The histone peptides used in ITC including H3 (1-19 aa), H3K4/9/14/18ac (1-26 aa), H4 (1-20 aa), and H4K5/8/12/16ac (1-20 aa), were commercially produced (Scilight-Peptide, Beijing). Titrations were performed at 20°C. Data were analyzed using Origin 7.0.

### Data availability

Raw RNA-seq and ChIP-seq data have been deposited in the Gene Expression Omnibus (GEO) database with the accession code GSE245273.

## Acknowledgements

This work was supported by the National Natural Science Foundation of China to X. J. He (32025003).

## Author contributions

X.-J.H. conceived and supervised the project. F.Q., Q.-Q.Z., J.-X.Z., and X.-J.H. designed experiments, analyzed data, and wrote the manuscript. Q.-Q.Z., F.Q., and J.- X.Z. performed experiments. D.-Y.Y. and Y.-N.S. performed bioinformatics analysis. L.L. and S.C. performed mass spectrometry analysis.

## Supplemental information

**Supplemental Figure 1.**
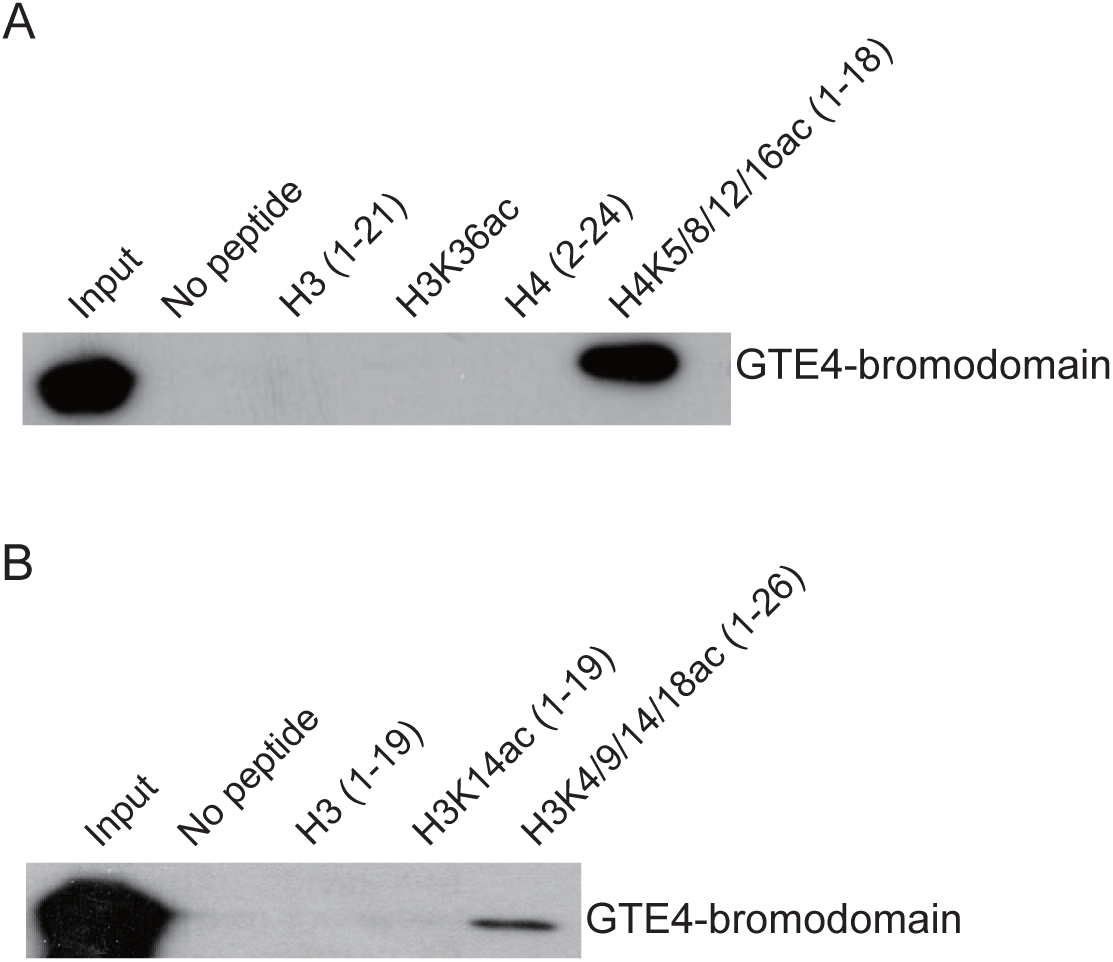
Determination of the binding ability of the bromodomain of GTE4 with acetylated histone by histone peptide pull-down assays. (A) The binding of the bromodomain of GTE4 to histone H4 with acetylation on lysine 5, 8, 12, and 16 (H4K5/8/12/16ac). (B) The binding of the bromodomain of GTE4 to histone H3 with acetylation on lysine 4, 9, 14, and 18 (H3K4/9/14/18ac). In the histone peptide pull-down assay, biotin-labelled histone peptides bound to the GST-tagged bromodomain of GTE4 were eluted by Streptavidin MagneSphere Paramagnetic Particles and detected by immunoblotting using anti-GST antibody.

**Supplemental Figure 2.**
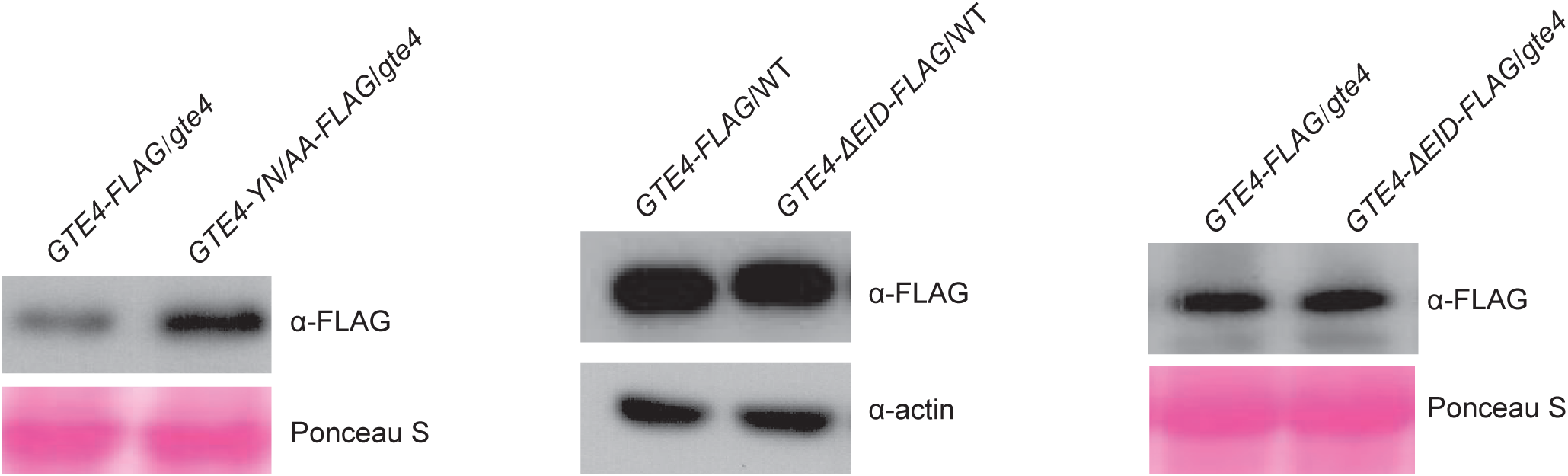
The expression levels of wild-type and mutated versions of FLAG-tagged *GTE4* transgenes. The expression levels of wild-type and mutated versions of *GTE4-FLAG* in the wild-type or *gte4* mutant backgrounds were determined by immunoblotting. The wild-type *GTE4-FLAG* and mutated *GTE4-YN/AA-FLAG* transgenic plants in the *gte4* mutant background were used for complementation testing and for the ChIP-seq analysis. The wild-type *GTE4-FLAG* and *GTE4-ΔEID-FLAG* transgenic plants in the wild-type Col-0 background were used for the AP-MS analysis. The wild-type *GTE4-FLAG* and *GTE4-ΔEID-FLAG* transgenic plants in the *gte4* mutant background were used for complementation testing and for the ChIP-seq analysis. The Ponceau S-stained protein or the actin protein detected by anti-actin antibody is shown as a loading control.

**Supplemental Figure 3.**
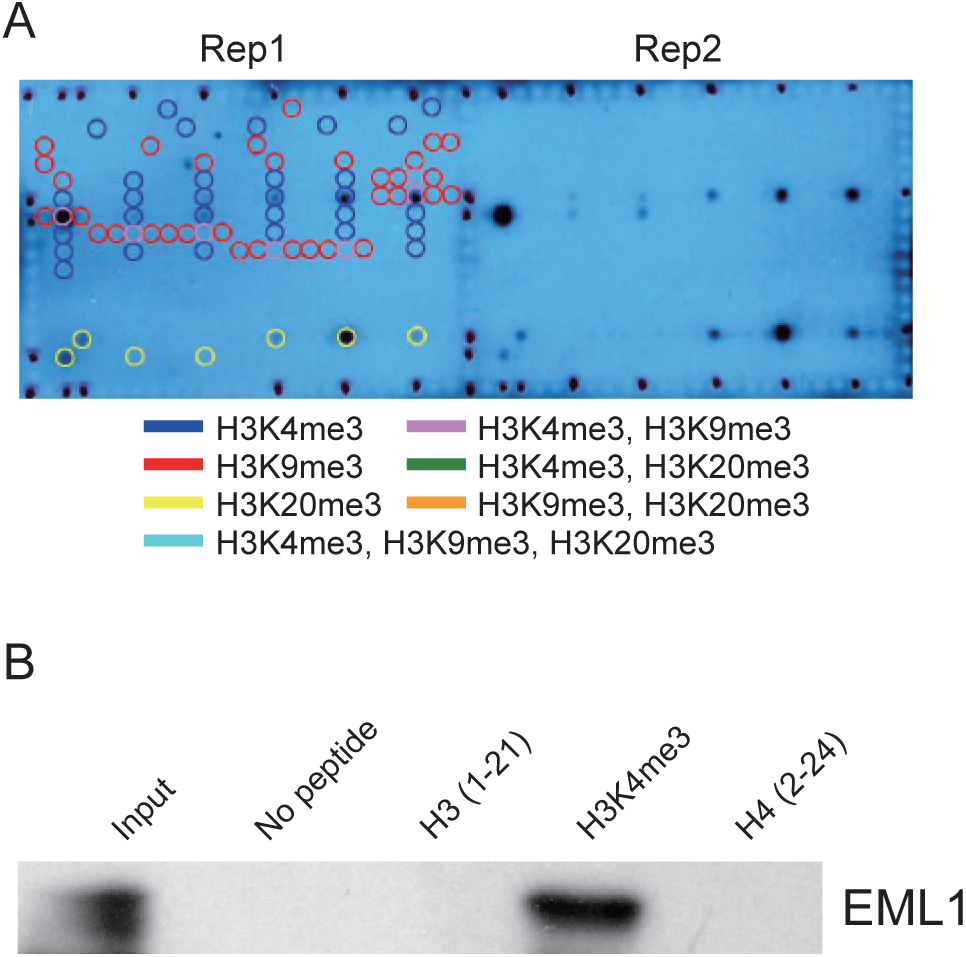
Identification of the EML1-recognized histone modification by a histone peptide array. (A) Images of the histone peptide array. The bacterially expressed GST-fused EML1 protein was loaded onto the histone peptide array and then subjected to the binding assay. The spots with indicated histone peptides were highlighted by different colors of circles. (B) Validation of the binding of EML1 to the H3K4me3 peptide by the histone peptide pull-down assay. The unmodified N-terminal peptides of H3 and H4 were used as negative controls.

**Supplemental Figure 4.**
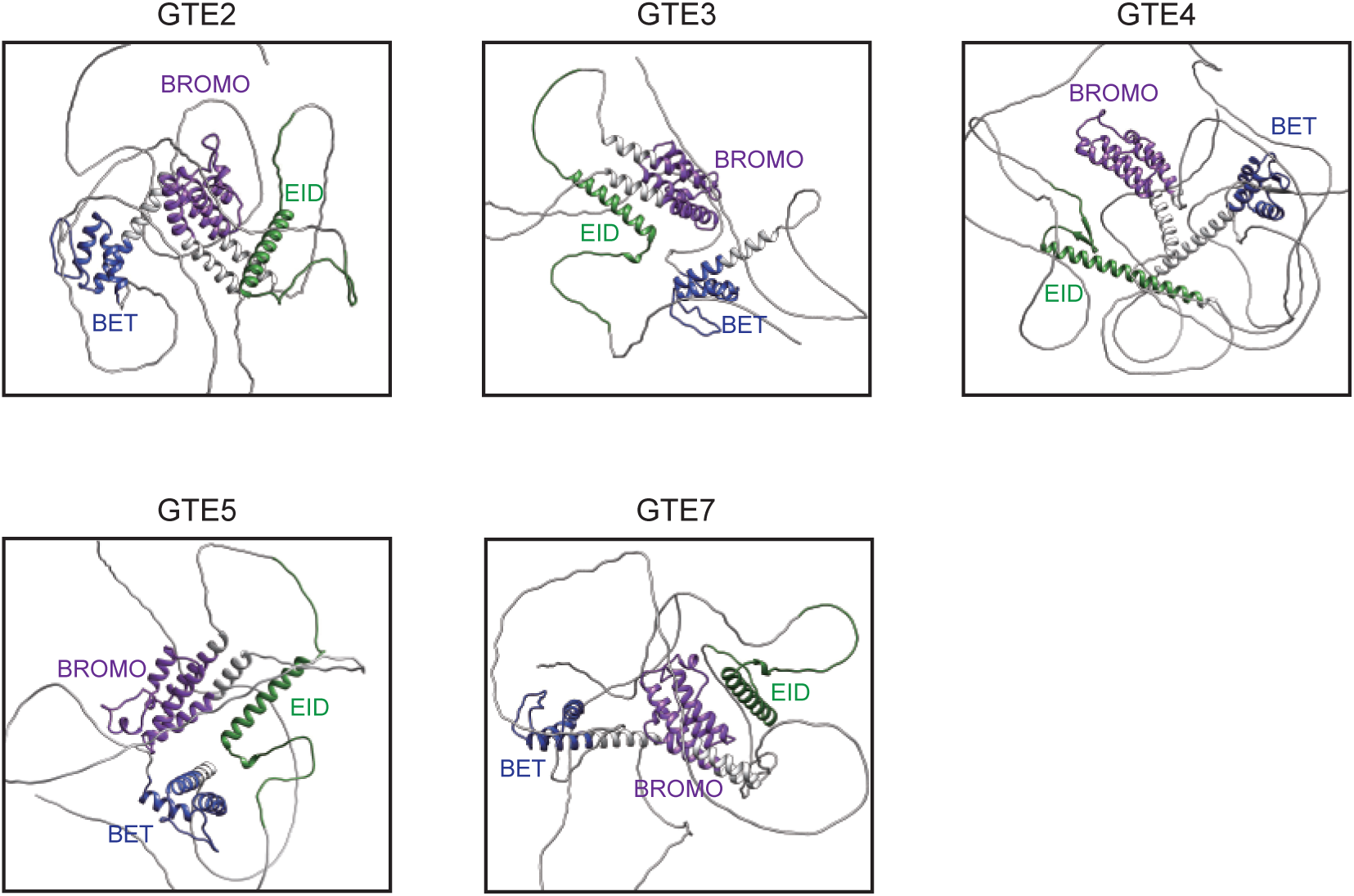
Structural prediction of GTE family proteins by AlphaFold. The structures of GTE2, GTE3, GTE4, GTE5, and GTE7, as determined by AlphaFold, are shown. The conserved EID, Bromo, and BET domains are indicated in green, purple, and blue, respectively.

**Supplemental Figure 5.**
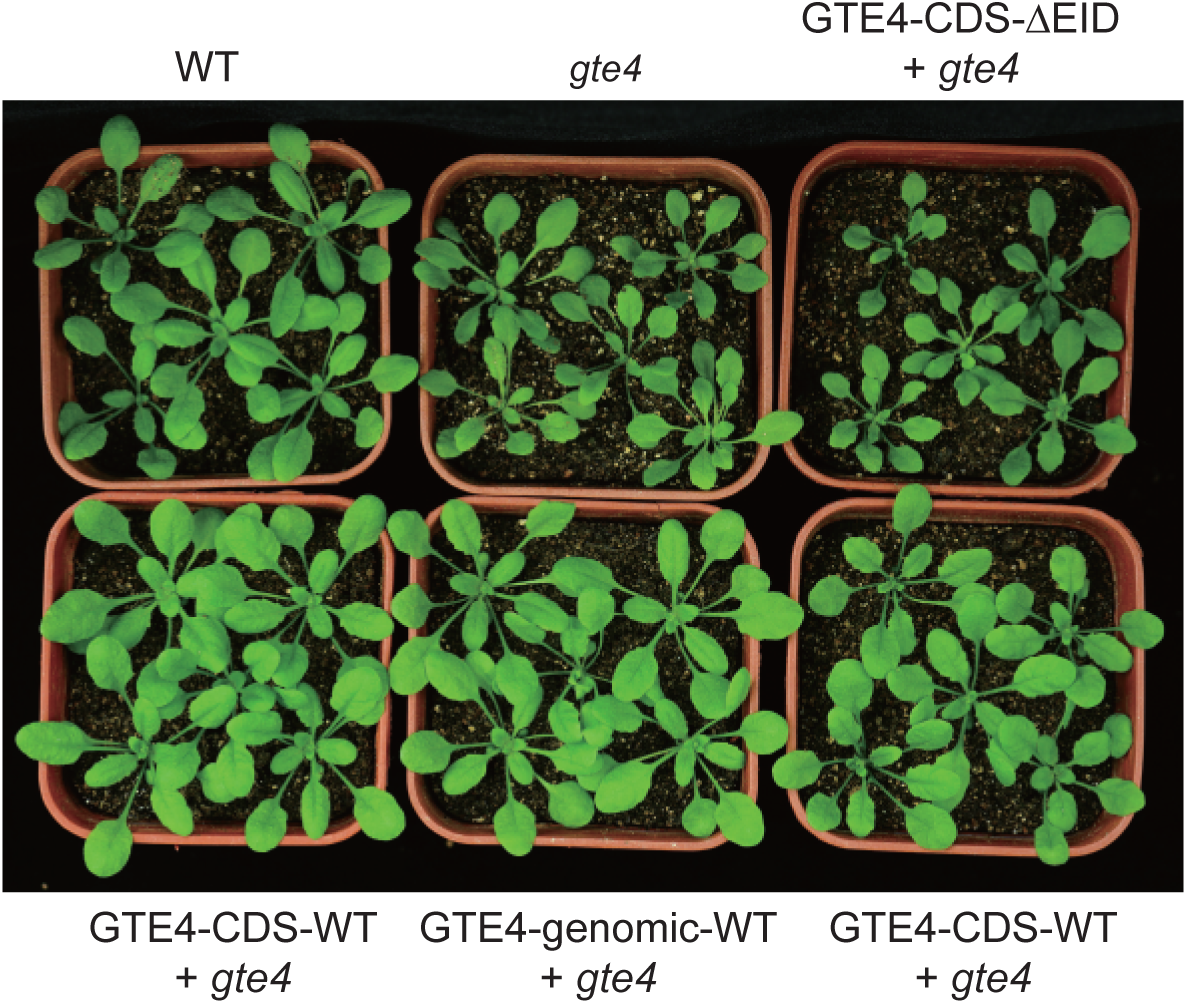
Determination of the impact of the EID deletion on the function of GTE4 by complementation testing. The wild-type and EID-deleted *GTE4* coding sequences driven by a native promoter were transformed into the *gte4* mutant for complementation testing. The wild-type *GTE4* genomic sequence driven by a native promoter was also transformed into the *gte4* mutant as a control. The image of 25-day-old plants with indicated genotypes is shown.

**Supplemental Figure 6.**
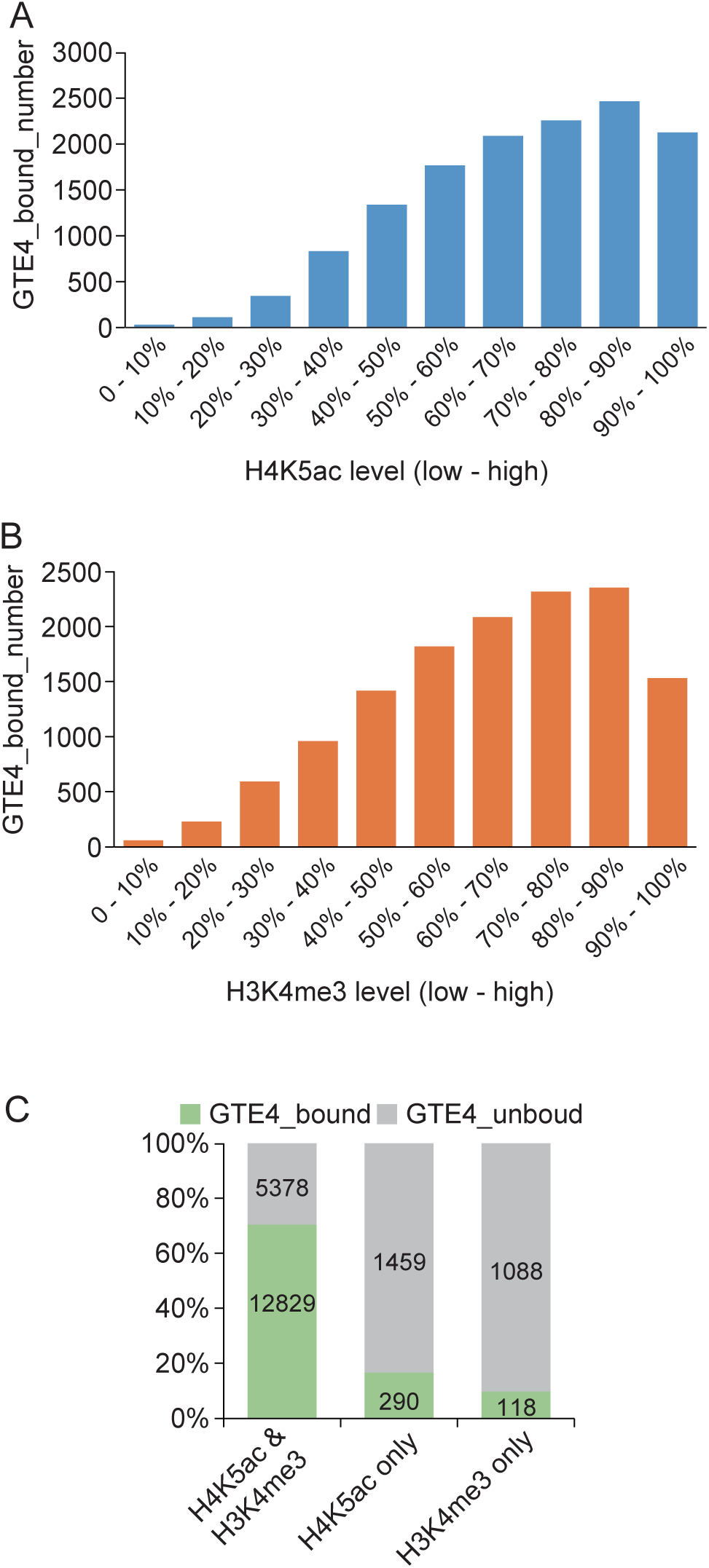
The effect of H4K5ac and H3K4me3 on the distribution of GTE4-bound genes at the whole-genome level. (A, B) The number of GTE4-bound genes within deciles of genes with incremental levels of H4K5ac (A) and H3K4me3 (B). (C) The ratio of GTE4-bound genes to genes co-occupied by H4K5ac and H3K4me3, genes exclusively occupied by H4K5ac, and genes exclusively occupied by H3K4me3.

**Supplemental Figure 7.**
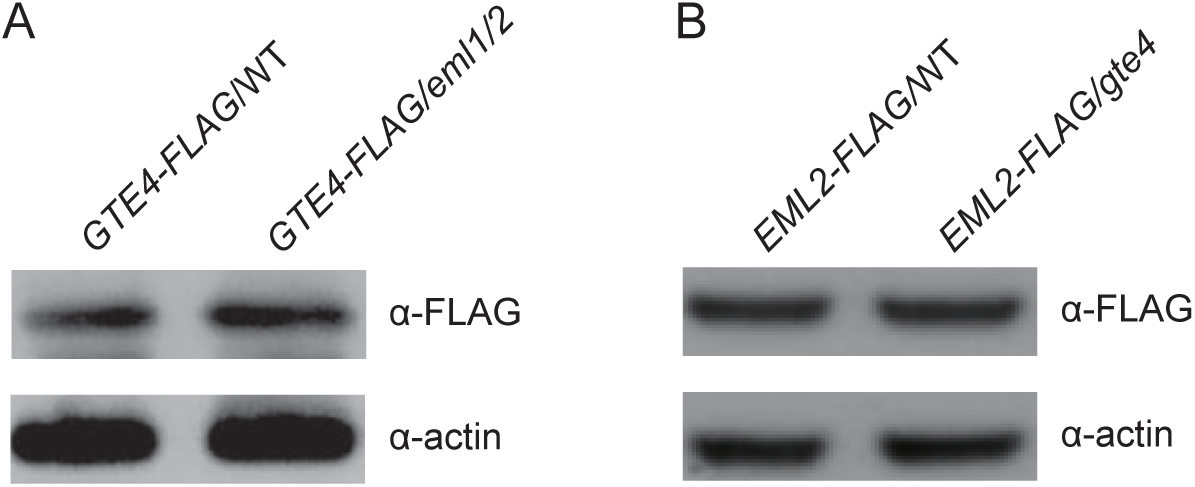
Determination of the expression levels of *GTE4-FLAG* and *EML2-FLAG* transgenes by immunoblotting. (A) The expression level of *GTE4-FLAG* transgene in the wild-type and *eml1/2* mutant plants. (B) The expression level of *EML2-FLAG* transgene in the wild-type and *gte4* mutant plants by immunoblotting. In (A) and (B), the expression levels of and *GTE4-FLAG* and *EML2-FLAG* were determined by immunoblotting using anti-FLAG antibody, and the actin signal was determined using anti-actin antibody and indicated as a loading control.

**Supplemental Dataset 1. Identification of acetylated histone-interacting proteins by AP-MS using histone H4K5/8/12/16ac peptides.**

**Supplemental Dataset 2. Identification of GTE4-, GTE4-ΔEID-, EML1-, and EML2-interacting proteins by AP-MS in Arabidopsis plants.**

**Supplemental Dataset 3. Differentially expressed genes identified by RNA-seq in *gte4*, *eml1/2*, and *gte4/eml1/2* relative to the wild type.**

**Supplemental Dataset 4. List of GTE4-enriched peaks and genes in the wild-type and *eml1/2* mutant backgrounds.**

**Supplemental Dataset 5. List of EML2-enriched peaks and genes in the wild-type and *gte4* mutant backgrounds.**

**Supplemental Dataset 6. H4K5ac and H3K4me3 levels of GTE4- and EML2-enriched genes.**

**Supplemental Dataset 7. Determination of the effect of *eml1/2* on GTE4 enrichment at GTE4-bound peaks and the effect of *gte4* on EML2 enrichment at EML2-bound peaks.**

**Supplemental Dataset 8. List of peaks and genes enriched by wild-type GTE4, GTE4-Y497A/N498A, and GTE4-ΔEID.**

**Supplemental Dataset 9. Determination of the effect of ΔEID and Y497A/N498A on GTE4 enrichment at GTE4-enriched peaks.**

**Supplemental Dataset 10. DNA oligos used in this study.**

